# Assessing the measurement transfer function of single-cell RNA sequencing

**DOI:** 10.1101/045450

**Authors:** Hannah R. Dueck, Rizi Ai, Adrian Camarena, Bo Ding, Reymundo Dominguez, Oleg V. Evgrafov, Jian-Bing Fan, Stephen A. Fisher, Jennifer S. Hernstein, Tae Kyung Kim, Jae Mun (Hugo) Kim, Ming-Yi Lin, Rui Liu, William J. Mack, Sean McGroty, Joseph Nguyen, Neeraj Salathia, Jamie Shallcross, Tade Souaiaia, Jennifer Spaethling, Chris P. Walker, Jinhui Wang, Kai Wang, Wei Wang, Andre Wilberg, Lina Zheng, Robert H. Chow, James Eberwine, James A. Knowles, Kun Zhang, Junhyoung Kim

## Abstract

Recently, measurement of RNA at single cell resolution has yielded surprising insights. Methods for single-cell RNA sequencing (scRNA-seq) have received considerable attention, but the broad reliability of single cell methods and the factors governing their performance are still poorly known. Here, we conducted a large-scale control experiment to assess the transfer function of three scRNA-seq methods and factors modulating the function. All three methods detected greater than 70% of the expected number of genes and had a 50% probability of detecting genes with abundance greater than 2 to 4 molecules. Despite the small number of molecules, sequencing depth significantly affected gene detection. While biases in detection and quantification were qualitatively similar across methods, the degree of bias differed, consistent with differences in molecular protocol. Measurement reliability increased with expression level for all methods and we conservatively estimate the measurement transfer functions to be linear above ~5-10 molecules. Based on these extensive control studies, we propose that RNA-seq of single cells has come of age, yielding quantitative biological information.

## Background

Single-cell RNA sequencing (scRNA-seq) allows unprecedented resolution for studies of gene expression. Since its introduction in 2009^1^, this approach has been used to identify and classify cell types, characterize rare cells, and study expression variation across cell populations^2–10^. In this method, the RNA content of a single cell is captured, reverse transcribed to generate cDNA, amplified and sequenced, providing measurements of the transcriptomes of single cells with nucleotide-level resolution. Compared with methods to sequence bulk RNA, scRNA-seq requires substantial molecular amplification and consequently, additional handling and enzymatic reactions. This has the potential to introduce additional experimental errors and molecular biases, such that analytic methods designed for bulk RNA sequencing may not be appropriate for single-cell measurements. Despite substantial experimental methods^11–16^ development, these measurements remain complex and poorly characterized. Though measurement characteristics likely depend on the specific experimental protocol used, there has been limited examination of whether measurement characteristics differ across methods. Additionally, although several common applications of single-cell RNA sequencing rely on measurement sensitivity, there are few assessments of the detection of gene expression and the factors that may affect it.

Here, we first describe a methodology to dissect the factors that affect scRNA-seq and then we characterize expression measurements generated by three scRNA-seq methods in terms of sensitivity, precision and accuracy. We find that all methods perform comparably overall, but that individual methods demonstrate unique strengths and biases.

## Results

### Method overview

Our approach was to dilute bulk total RNA (from a single source) to levels bracketing single-cell levels of total RNA (10 pg and 100 pg). Here, we analyzed the performance of scRNA-seq methods in terms of sensitivity (number of unique gene models detected), precision (replicate variation), and accuracy (deviation from bulk). We note that expected replicate precision depends on the exact sequence of dilutions that lead to the final set of replicates. For example, if a single “master” dilution mix is made from which *n* replicate final dilutions are created, the expected number of molecules for each replicate will be based on the master dilution, not the original bulk. Each replicate value in relation to the bulk will be comprised of two terms, the variance term due to the final dilutions and a bias term, which is the deviation of the master dilution from the bulk. Different experimental protocols (e.g., using Fluidigm C1 to generate replicates) require attention to the expected variation. Overall, we adopted the framework of estimating a “transfer function”—i.e., the function that describes the input-output relationship of an instrument. As reported below, various factors affect the transfer function and we employed a general linear model framework to dissect the factors governing measurement performance. In particular, we observed that certain genes, or even control ERCC probes, have a tendency for large deviations from expectations and we created a list of problematic gene models for future reference. We add the caveat that our transfer functions are not reliable outside the range of experimental values from which we fitted the models and the inferences should be interpreted with care.

### RNA-sequencing datasets

We performed replicate transcriptome amplifications of Universal Human Reference RNA (UHR) and Human Brain Reference RNA (HBR) that were diluted to single-cell and ten-cell abundances (10 and 100 picograms (pg.) total RNA or ~200,000 and 2 million mRNA molecules, respectively) and were amplified using three single-cell RNA amplification methods (Figure 1A-B). Methods included the antisense RNA IVT protocol (aRNA), a custom C1 SMARTer protocol (SmartSeq Plus) performed on a Fluidigm C1 96-well chip, and a modified NuGen Ovation RNA sequencing protocol (NuGen, Figure 1B-C, Table S1). Bulk ribo-depleted UHR and HBR RNA were sequenced and served as a reference. The general experimental scheme was consistent for all dilution replicates; however, there were differences across experimental groups in the specifics of experimental protocols, necessitated by particular methodologies (Figure 1A, see *Methods* and Table S1 for full details). Because of these experimental differences, head-to-head comparison of methods is not appropriate and our goal is to provide quantitative analyses of factors affecting individual methods. Current results should be used in experimental planning, data analysis, and method optimization rather than as a performance test of any particular method.

**Figure 1.**
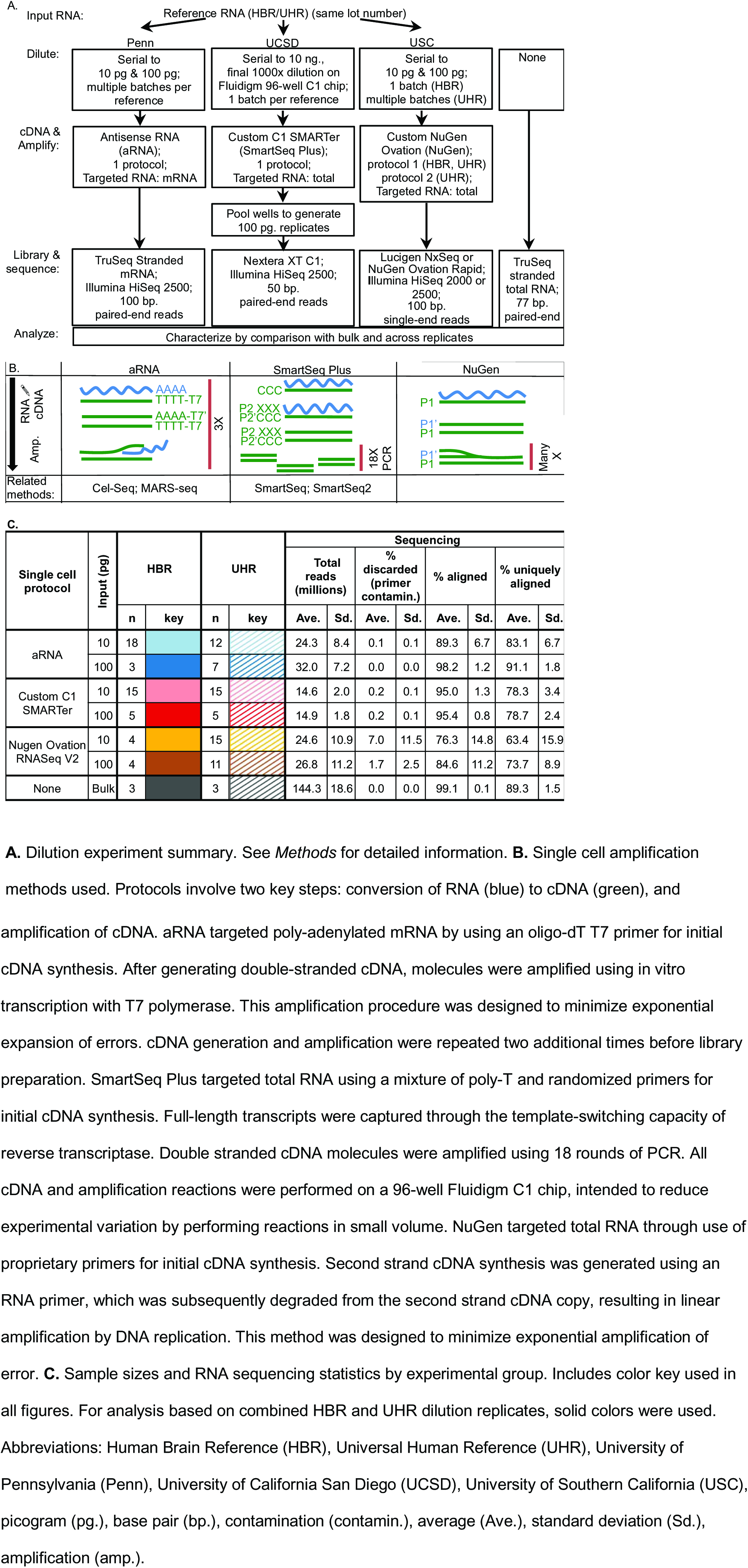
Experimental design and RNA sequencing statistics by experimental group

### Data processing

Briefly, all samples were aligned to hg19 using STAR aligner^17^. Uniquely aligned reads were assigned to GENCODE18 gene annotations using HTSeq and htseq-counts^18^ and then were depth normalized^19^. Ribosomal genes and genes with short isoforms (<300 nucleotides) were excluded because of differences in sequencing protocols across groups (Figure 1A), leaving 42,855 genes for analysis. (We use “gene” to match GENCODE18 gene ids, a set that includes both coding‐ and non-coding RNA.) T o avoid artifacts caused by alignment or quantification ambiguities, we generated a stringently filtered gene list containing 10,039 genes to which reads can be uniquely assigned and referred to these genes as “computationally unambiguous” throughout (Table S2). Reference RNA were aligned and quantified with RSEM (RNA-seq by Expectation-Maximization)^20^. Estimated abundances were concordant with publicly available PrimePCR measurements and with poly-A RNA sequencing measurements (Figure S1, SEQC/MAQC-III Consortium, 2014, GEO accession numbers: GPL18522, GSM1362002-GSM1362029, GSM1361974-GSM1362001^21^). The mass of targeted input RNA in diluted replicates was estimated as in Brennecke et al.^2^ and was used to calculate, for each gene, the expected number of input molecules in a diluted replicate. aRNA selectively targeted poly-adenylated (poly-A) mRNA (Figure 1A). We calculated the expected number of input poly-A molecules using publicly available bulk HBR sequencing measurements. See *Methods* for further details.

On average, replicates were sequenced at a depth of 22.0 ± 9.6 million reads (± standard deviation or Sd.). 1.5 ± 5.3% of reads were discarded due to primer contamination. 89.3 ± 10.6% of retained reads aligned to the genome, 77.6 ± 11.2% uniquely (Figure 1C). To examine the coverage distribution of each method, we quantified the frequency of mapped reads over several genomic regions of interest (Table 1). This distribution differed for the three single-cell amplification methods. The majority of aligned reads for aRNA dilution replicates originated from non-mitochondrial exons (excluding rRNA), a substantially larger proportion than that recovered by SmartSeq Plus or NuGen. rRNA genes, pseudogenes and repeats encoded by the nuclear genome comprised a small fraction of reads in all amplified libraries (average ± SD: 0.67 ± 0.65%). rRNA and mRNA encoded by the mitochondrial genome (2 genes and 13 genes, respectively) constituted a substantial percentage of reads (average ± SD: 16.5 ± 8.4%). Mitochondrial recovery differed substantially across methods. This difference may translate into a method-specific effect on depth normalization and for this reason mitochondrial genes have been excluded from the subsequent analyses. The distribution of reads across genomic features also differed substantially across replicates for aRNA and NuGen (Table 1).

**Table 1.** Coverage selectivity by method

| Source | Protocol | Genome coverage |  |  |  |  |  |  |  |  |  |
| --- | --- | --- | --- | --- | --- | --- | --- | --- | --- | --- | --- |
|  |  | % exons<br>(excluding rRNA<br>& mitochondria) |  | % intronic |  | % rRNA (nuclear) |  | % rRNA<br>(mitochondrial) |  | % mitochondrial<br>(non-rRNA) |  |
|  |  | Ave. | Sd. | Ave. | Sd. | Ave. | Sd. | Ave. | Sd. | Ave. | Sd. |
| HBR | aRNA | 59.07 | 5.70 | 23.29 | 4.68 | 0.03 | 0.02 | 4.82 | 1.62 | 12.29 | 2.79 |
|  | SmartSeq Plus | 41.00 | 0.86 | 39.56 | 0.85 | 1.28 | 0.07 | 10.02 | 0.43 | 7.77 | 0.26 |
|  | NuGen | 29.27 | 4.97 | 45.09 | 5.88 | 1.33 | 0.40 | 20.38 | 5.19 | 3.65 | 0.59 |
|  | Bulk (Poly-A) | 80.13 | - | 8.52 | - | 0.08 | - | 1.96 | - | 9.01 | - |
|  | Bulk (rRNA-depleted) | 61.44 | 0.54 | 37.89 | 0.45 | 0.03 | 0.01 | 0.10 | 0.04 | 0.25 | 0.04 |
| UHR | aRNA | 71.57 | 2.34 | 23.02 | 2.60 | 0.06 | 0.05 | 1.61 | 0.19 | 3.38 | 0.59 |
|  | SmartSeq Plus | 35.89 | 0.89 | 52.84 | 0.94 | 0.31 | 0.02 | 6.20 | 0.26 | 3.85 | 0.12 |
|  | NuGen | 33.00 | 4.02 | 39.33 | 8.99 | 1.25 | 0.57 | 23.47 | 7.31 | 2.59 | 0.62 |
|  | Bulk (Poly-A) | 86.99 | - | 7.11 | - | 0.11 | - | 0.47 | - | 5.09 | - |
|  | Bulk (rRNA-depleted) | 58.17 | 0.34 | 41.28 | 0.29 | 0.02 | 0.01 | 0.03 | 0.01 | 0.18 | 0.02 |

Average percent of aligned reads assigned to genomic regions for each method. Nuclear rRNA includes rRNA genes, pseudogenes and repeats. See *Methods* for definitions of genomic regions.

### Gene detection sensitivity

We calculated the number of detected genes as a measure of detection sensitivity (Figure 2A). All methods demonstrate comparable high gene detection, detecting greater than 70% of the expected number of genes, with SmartSeq Plus demonstrating the highest detection (Byar’s 95% C.I., Obs. / Exp.: aRNA (0.722, 0.726); SmartSeq Plus (0.877, 0.882); NuGen (0.735, 0.740)). With respect to poly-A RNA, aRNA detected (0.840, 0.844) of expectation. Variation across samples within each method was substantially larger than expected due to dilution suggesting additional loss during cDNA and amplification (Figure 2A).

**Figure 2.**
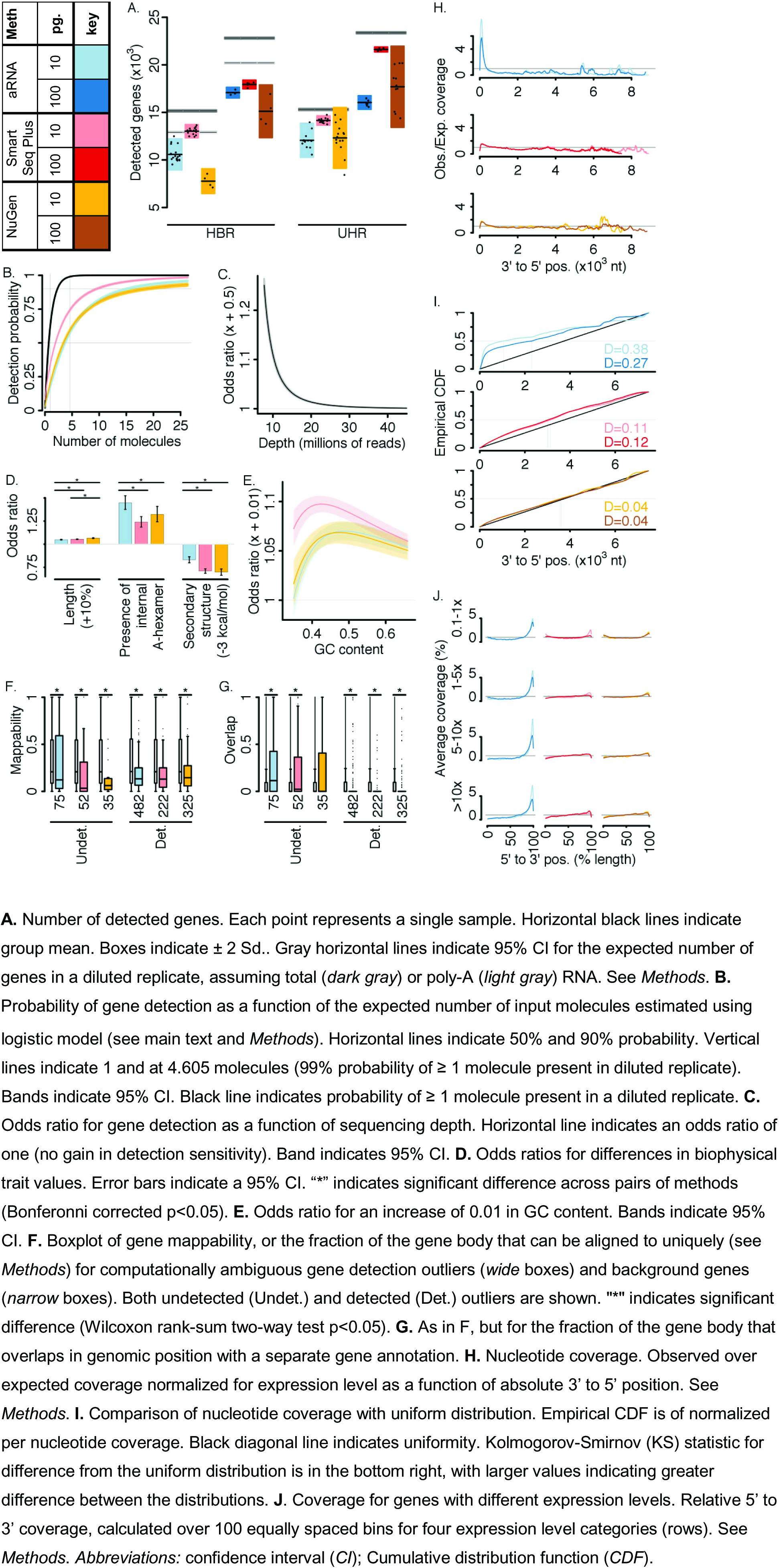
Single-cell RNA sequencing sensitivity

Detection of a given gene may depend on parameters such as the input number of molecules, GC-content, presence of internal adenosine monophosphate (A) hexamers, length, strength of molecular secondary structure, and sequencing depth. To estimate the contribution of these factors to gene detection, we fit a logistic regression model to the 10 pg. gene detection data with gene detection as the dependent variable, considering only computationally unambiguous genes to focus on experimental sensitivity. (See *Methods* and Tables S3-S4 for details.) All methods had a 50% probability of gene detection at ~2-4 expected input molecules, controlling for the remaining covariates (Figure 2B, Table S5). We calculated a molecular recovery rate as the predicted probability that a gene with 1 expected input molecule will be detected, scaled by the probability that at least one molecule of such a gene will be in a diluted replicate. Molecular recovery rates were greater than 0.25 for all methods (95% prediction interval: aRNA (0.262, 0.279), SmartSeq Plus (0.534, 0.558), NuGen (0.315, 0.339)). With respect to poly-A RNA, aRNA recovery rate was (0.320, 0.349).

Despite the small number of total (targeted) RNA molecules in a single 10 pg. dilution replicate (estimated here to be ~300,000 molecules), sequencing depth had a highly significant effect on gene detection (Table S3). Figure 2C shows the odds ratio of increasing sequencing depth by 500,000 reads. The odds ratio is the relative odds of event occurrence at two values of a variate, controlling for all other covariates. The odds of gene detection increased substantially with sequencing depth until a depth of ~15-20 million reads or ~50 reads per input molecule. Here, increasing sequencing depth from 10 to 15 million reads translated into an expected gain of 25.02 % in detected genes. The influence of remaining covariates on gene detection differed across methods (Figure 2D-E). The odds of gene detection increased with gene length, and NuGen demonstrated a significantly stronger length effect than aRNA or SmartSeq Plus (Figure 2D). The presence of an internal A-hexamer positively influenced the probability of gene detection for all methods, with strongest effect for aRNA. Increased strength of secondary structure decreased the odds of detection for all methods, with significantly smaller effect for aRNA than for SmartSeq Plus or NuGen. While GC content influenced detection probability in a complex manner, SmartSeq Plus demonstrated the strongest GC effect (Figure 2E).

A small fraction of computationally unambiguous genes had poor fit by the logistic model (0.30 ± 0.14%; see Table S6 for a list of outliers and *Methods* for details). Each outlier was categorized as “detected” if the gene was unexpectedly observed and “undetected” if it was unexpectedly missing.

Nearly all identified outliers (16/17) were method-specific. A larger proportion of computationally ambiguous genes were poorly fit by the model (3.21 ± 0.23 %, Table S6) with a sizable fraction (19.81 ± 2.90%) that fit poorly for all methods. These outlier genes had significantly lower fraction of the gene body that could be aligned uniquely than background genes (Figure 2F; Wilcoxon rank sum two-way test, p<0.05). This was the case for both detected and undetected outliers, indicating that alignment ambiguities likely generate both false positives and false negatives. Outliers also significantly differed from background in the fraction of the gene body that overlaps with another gene annotation, with lower overlap among detected outliers and greater overlap among undetected outliers (Figure 2G).

To characterize read coverage at the scale of individual base positions, we calculated the observed / expected nucleotide coverage as a function of 3' to 5' position within a gene (Figure 2H), normalized such that a uniform distribution of reads along a gene would be assigned a value of one at all positions (see *Methods).* Coverage for all methods was significantly different from uniform (Figure 2I; Kolmogorov-Smirnov test p<10^-10^ for all groups); however, NuGen demonstrated the greatest uniformity (Figure 2H-I) with similar positional coverage distribution for 10 pg. and 100 pg. dilution replicates. aRNA preferentially covered the 3’ terminal and demonstrated greater 3’ bias for 10 pg. dilution data. SmartSeq Plus showed an intermediate degree of bias. Segregated by expression levels, we found preferential recovery of the 5’ and the 3’ gene ends for low abundance genes and preferential 3’ coverage for high abundance genes (Figure 2J).

### Precision

We next consider the similarity of measurements across dilution replicates, within methods and across methods. Though we cannot quantitatively compare measurement precision across methods (see Figure 1A), the results will be applicable to experimental design and analysis for each method. The average within experimental group pairwise correlation coefficient (± Sd.) was 0.37 ± 0.07 (Kendall) and 0. 51 ± 0.09 (Pearson, log_10_ counts) for 10 pg. replicates and 0.64 ± 0.06 (Kendall) and 0.79 ± 0.06 (Pearson, log_10_ counts) for 100 pg. replicates (Figure 3A; zeros treated as missing values).

**Figure 3.**
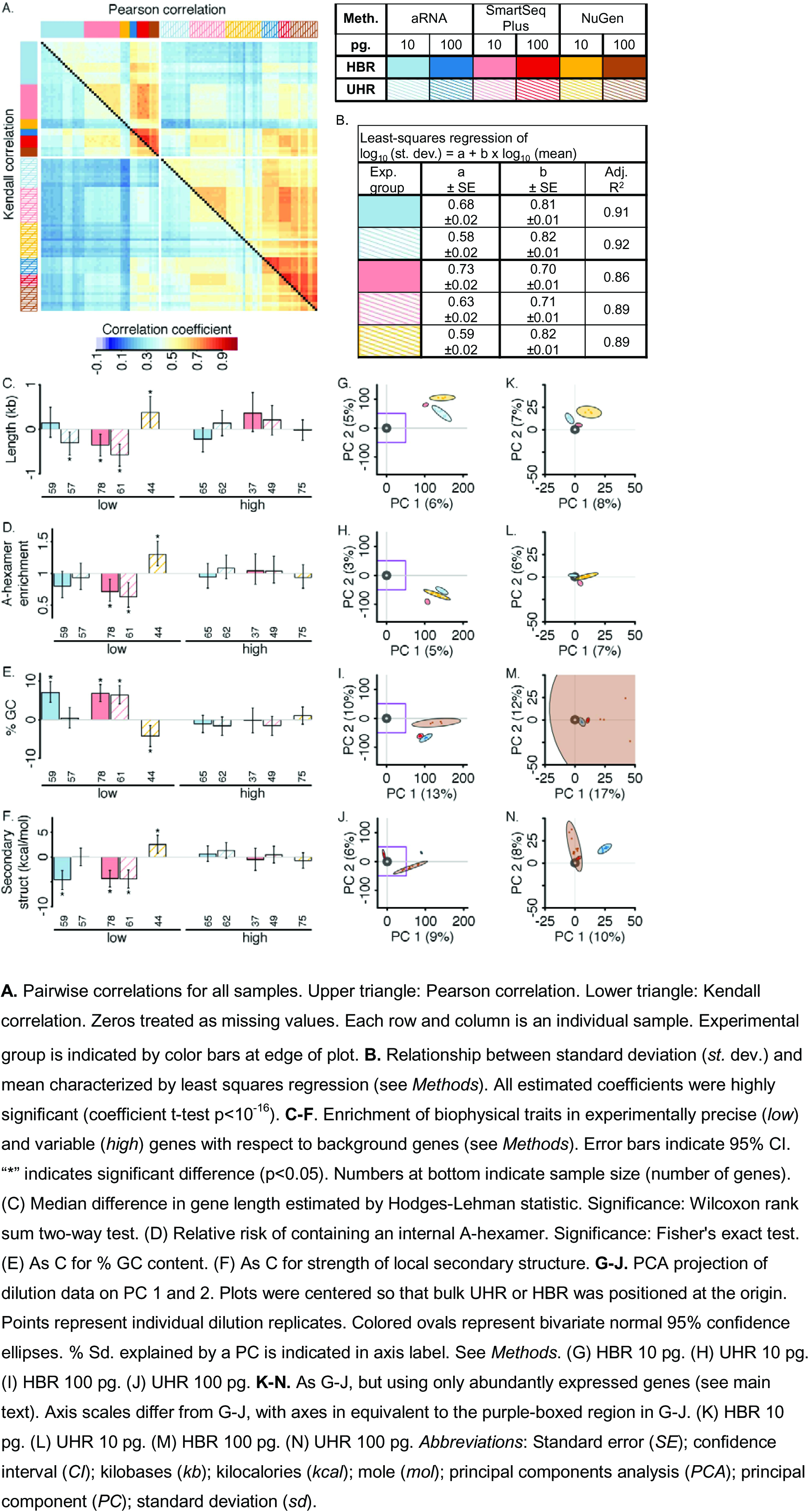
Single-cell RNA-sequencing precision

To describe the dependence of precision on expression level, we performed least-squares regression of the empirical standard deviation on the empirical mean (both variables log-transformed to satisfy the assumption of residual normality) for 10 pg. experimental groups with sample size >5. The mean was an excellent predictor of standard deviation (Figure 3B, adjusted R^2^>0.85 and slope coefficient t-test p <10^-16^ in all cases). Genes that were less precise than expected (standardized residuals outside predicted 90% CI) differed little from background in their biophysical characteristics (Figure 3C-F), suggesting limited systematic bias in experimental variability. Biophysical characteristics enriched among unexpectedly precise genes with respect to background differed in a method-specific manner (Figure 3C-F). For aRNA and SmartSeq Plus, enriched biophysical characteristics were concordant with reduced probability of gene detection (compare Figure 2D-E), suggesting technical dropouts might play a strong role in replicate precision. NuGen demonstrated the opposite trend suggesting that amplification bias might play a stronger role. A subset of genes whose standard deviation was poorly predicted by the mean (percent of genes with standardized residuals outside predicted 99.3% C.I.) are listed in Table S7. We recommend that the expression values of these gene models should be interpreted with caution.

Separate principal components analysis (PCA) of each HBR and UHR for 10 pg. dilution data demonstrated that average displacement between single cell and bulk measurements predominate over differences between single cell methods (Figure 3G-H); however, there were clear differences across methods in the multivariate covariance structure of experimental variation. Differences across methods were also apparent for 100 pg. dilution replicates (Figure 3I-J), and, though these measurements were more similar to bulk measurements, differences between dilution replicates and bulk measurements persisted. We note that average displacement between single cell and bulk measurements represent both a bias component from utilizing a master dilution mix (see above) and technical bias. We repeated PCA on a subset of genes with greater than 18.5 expected input molecules (expected probability of detection for “typical” gene > 0.9 for all methods). On highly abundant genes, dilution replicates were substantially more similar to bulk measurements (Figure 3K-N) and differences across methods were substantially smaller. However, in all cases, the within method pattern of covariation (direction of ellipses) and the bias dispersal around the bulk expected value (position of the centroid of the ellipses) differed for both source RNA and individual methods.

### Accuracy

We calculated pairwise correlation coefficients of dilution replicates with bulk as a metric of overall accuracy (Figure 4A). For this and the below, only non-zero gene counts were considered in order to focus on quantitation rather than sensitivity. 10 pg. dilution replicates demonstrated an average pairwise correlation with reference of 0.42 ± 0.01 (Kendall) and 0.55 ± 0.01 (Pearson, log_10_ counts). 100 pg. replicates showed greater similarity with reference (0.57 ± 0.01 (Kendall) and 0.72 ± 0.01 (Pearson, log_10_ counts). Correlation with reference had a modest association with percent unique alignment (Figure 4B-C).

**Figure 4:**
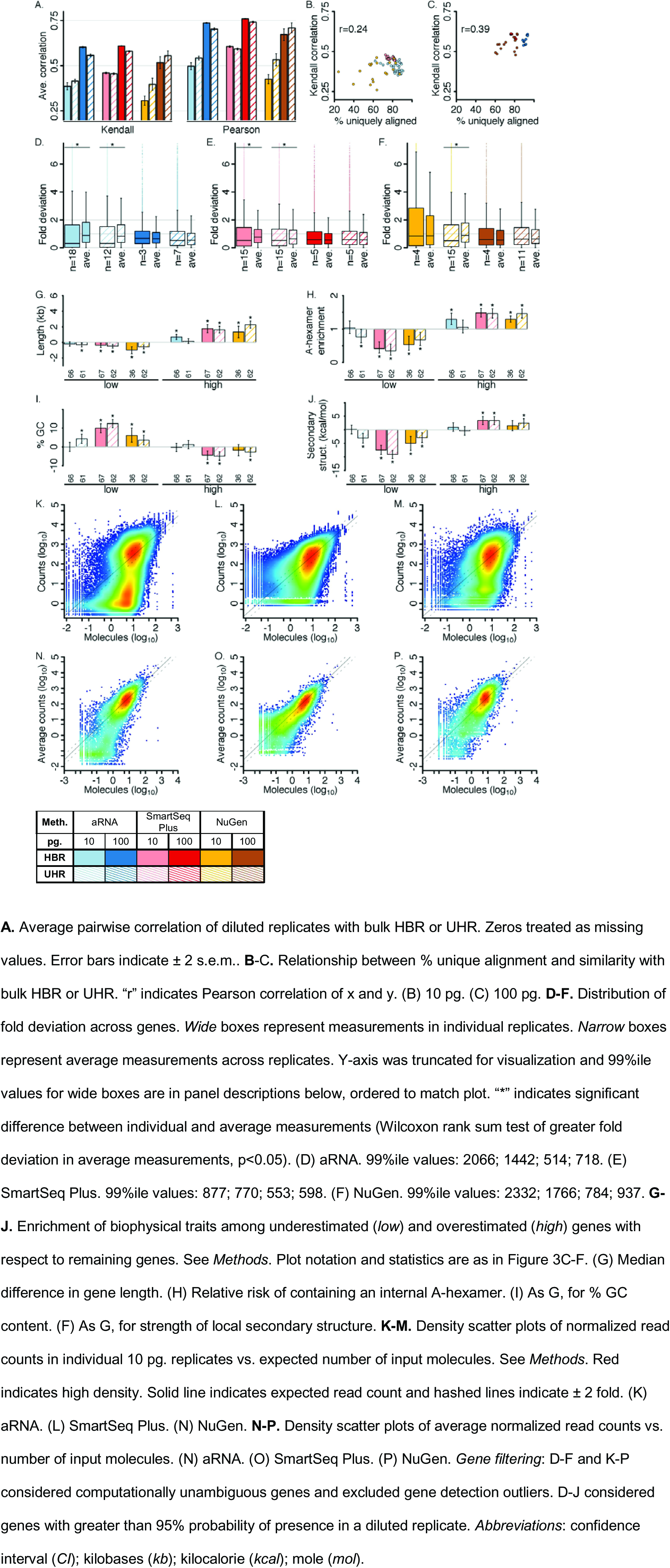
Single-cell RNA sequencing accuracy

To assess the accuracy of individual gene estimates, we calculated the fold deviation of normalized read counts with respect to bulk HBR or UHR measurements (Figure 4D-F, *Methods*). For all methods and input amounts, the median fold deviation was less than 1 but a subset of genes was extensively overestimated. Overestimated genes (top 5% fold-deviation) were substantially longer than remaining genes and more frequently contained an internal A-hexamer (Figure 4G-J). For NuGen and SmartSeq Plus, these genes also had lower GC content and weaker local secondary structure than remaining genes. Underestimated genes (bottom 5% fold-deviation) demonstrated the opposite tendencies: compared to background genes, they were shorter, less frequently contained internal A-hexamers, had higher GC content and stronger secondary structure than background, as might be expected (Figure 4G-J). Overall, aRNA demonstrated less systematic bias than NuGen or SmartSeq Plus. Highly inaccurate genes (top or bottom 1% fold-deviation) are catalogued in Table S8.

Smoothed density scatter plots demonstrated method-specific transfer functions between the expected number of input molecules and the number of read counts in an individual replicate (Figure 4KM). This relationship was roughly linear at expression levels greater than ~5-10 expected input molecules up to at least ~600 input molecules, the highest expression level examined for 10 pg. replicates, giving a linear dynamic range of at least 100-fold. At low to mid expression levels measurements were frequently underestimated expanding the apparent range of measured abundances, particularly for aRNA and NuGen (Figure 4N-P).

### Protocol variations

We evaluated the effects of several protocol variations on measurement quality (Table 2). The aRNA protocol used for the primary analysis includes cDNA purification before initial amplification, and 3 rounds of IVT amplification followed by dilution of amplified cDNA before library preparation (Figure 1B). Elimination of initial cDNA purification significantly improved sensitivity and accuracy, as did reduction to two rounds of IVT amplification and elimination of dilution prior to library generation (Table 2). An optimized protocol incorporating both changes, demonstrated substantial improvements in the number of detected genes and pairwise correlation with the bulk (Table 2).

**Table 2.**
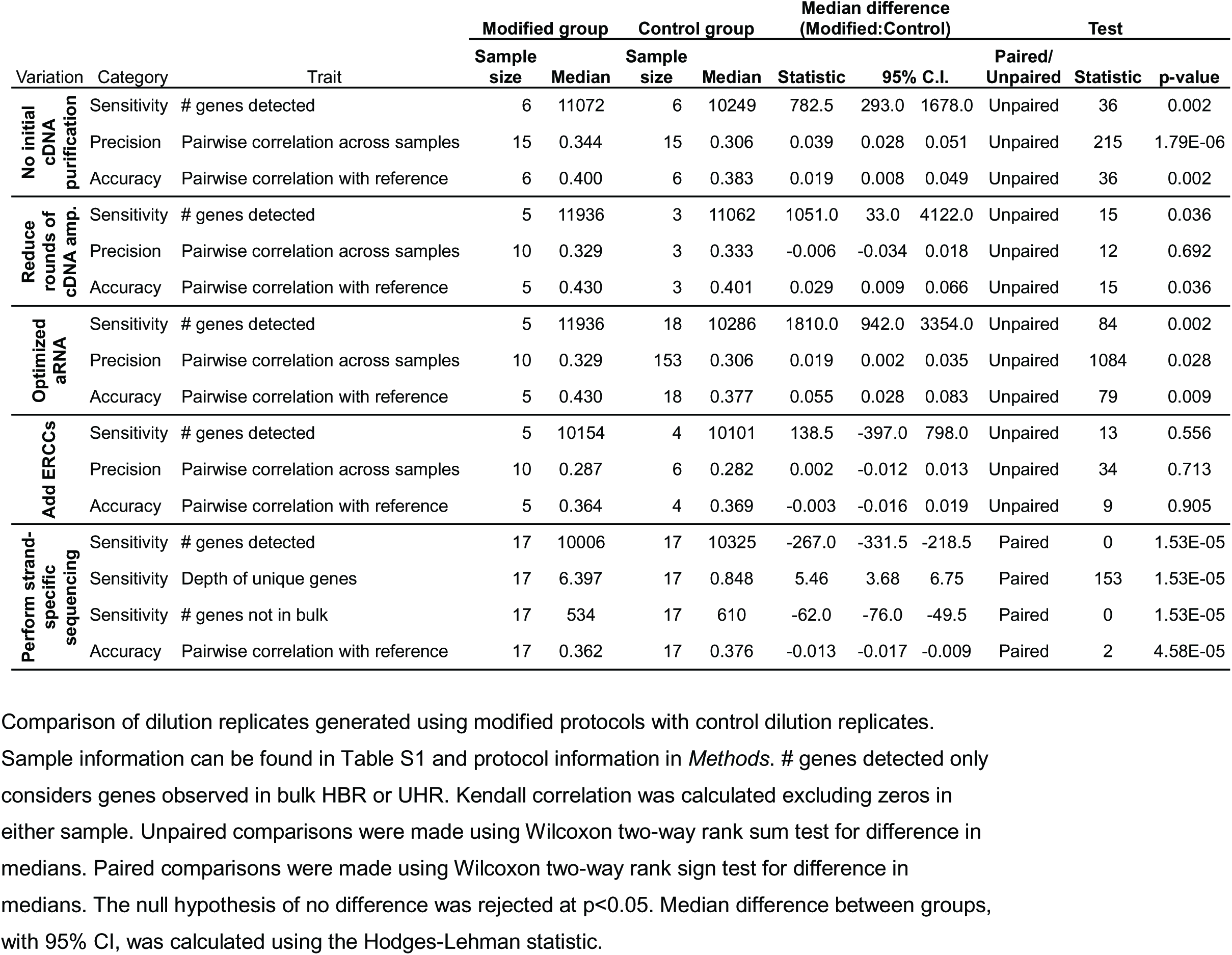
Evaluation of protocol variations

The addition of ERCC spike-in transcripts provides an internal control^22^, but it raises the concern that addition of synthetic RNA to a sample may decrease biological sensitivity. We found no significant difference in sensitivity, precision or accuracy across matched dilution replicates with and without the addition of ERCCs (Table 2) up to the spike-in level of 2.7% of reads. Individual ERCC transcripts were found to be problematic, consistently inaccurate, for SmartSeq Plus and aRNA in a method specific manner (Figure S2).

Strand-specific RNA sequencing may improve detection sensitivity and reduce false positive detection. Stranded quantification of aRNA replicates detected slightly fewer genes than non-stranded quantification; however, it also detected significantly fewer genes that were not observed in the bulk, and genes that were detected only by stranded quantification were supported by significantly more reads than genes detected only by non-stranded quantification (Table 2).

## Discussion

In light of these results, we briefly discuss a few topics related to experimental planning, method optimization and data analysis.

Though the goal of this study and our experimental design is not meant to select “the best method”, some results may be helpful in selecting an appropriate method for a particular project. The enriched coverage of exons in aRNA may be beneficial for studies of mRNA, and the retention of transcript strand information is unique to aRNA at this point. SmartSeq Plus and C1 microfluidic device generates reproducible replicates and high detection sensitivity, presumably due to more uniform liquid handling and retention of material due to lack of vessel transfer. The uniformity of coverage provided by NuGen (Figure 2H-I) may be beneficial studies of isoform use and splicing. We note that, in our hands, NuGen reactions were inconsistent and we had repeated amplification failures, or amplification of nontemplate directed products with this method, especially at the 10 pg level where the method appears to be reaching the limits of it sensitivity.

In selecting sequencing depth, there is a trade-off between gene detection sensitivity and cost. Typically, a small number of genes comprise the bulk of RNA molecules in a transcriptome. Sequencing at low depths should be sufficient to reproducibly detect and quantify these abundant genes; however, the majority of genes in a typical transcriptome are at low abundance. Because of this, the number of genes detected in the mixtures of RNAs used here depends heavily on sequencing depth (Figure 2C). The dynamic range limit due to sequencing depth is will be a function of the relative frequency distribution, which will vary for an actual single cell. Our results suggest that increasing the number of reads per cell may produce richer transcriptome measurements and should be considered carefully in the context of a specific experimental plan.

Missing values due to lack of sensitivity and the presence of large valued outliers may cause complications for depth normalization methods. Large variation across samples and substantial differences across methods in the fraction of reads assigned to mitochondrial RNA (Table 1) will propagate to sample and method differences in relative read counts. More generally, we observed large variation in the distribution of reads across broad genome annotation classes (Table 1). Because each genomic annotation class accounted for a substantial number of reads and input molecules, the observed differences across methods, and within methods, cannot be simply explained by sampling error. Similarly, variation across samples in the number of detected genes cannot be easily explained by dilution (Figure 2A). This behavior might be explained by global differences in reaction efficiencies across samples, as suggested previously^25^; however, the experimental sources of such differences in a controlled experiment are unclear. We found certain subsets of genes to be problematic for gene detection, accuracy, and precision, in a method-specific manner (Table S6-S8). We recommend that genes on these lists be treated with caution, filtered before analysis or interpreted with care. We similarly found several ERCC spike-in transcripts to be problematic (Figure S2), and recommend selecting a subset of reliable ERCC transcripts for use as reference measurements.

Some scRNA-seq quantification challenges might be reduced through further experimental optimizations, for example by increasing detection sensitivity and reducing amplification biases. Eliminating the initial cDNA purification, reducing the extent of amplification required, and limiting sample dilution may be productive avenues, as suggested by our data. Methods to experimentally deplete highly abundant and variably recovered mitochondrial RNA, if not of experimental interest, may also be of use.

Single cell RNA measurement methods have become increasingly robust and automated systems have made the technique broadly more accessible and efficient. All methods examined here demonstrated good gene detection and a linear relationship between input molecular abundances and measured expression levels at mid- to high-expression levels or greater than ~5-10 input molecules. This corresponds to ~4,000-8,000 reliably measured genes for the reference transcriptomes examined here. We propose that single cell RNA measurements have come of age and this level of resolution for gene expression measurements has and will continue to facilitate biological discovery.

## Methods

### Experimental design

Each collaborating center obtained reference RNA with the same lot number for Universal Human Reference (UHR) RNA (Agilent 740000, Lot 0006141415) and Human Brain Reference (HBR) (Ambion AM6050, Lot-105P055201A) and performed replicate amplification using a single amplification method, detailed below.

#### SmartSeq Plus

Reference RNA was diluted to an intermediate stock solution by serial dilution. A final 1000-fold dilution occurred on the C1 chip, such that individual wells in a given batch contained 9.99 pg. sampled from a common intermediate dilution. ERCC spike-in RNA mix 1 (Ambion 4456740) was also added for a final mass of approximately 7 femtograms (fg.) per sample, a 4,000,000x dilution from stock. Samples for each source RNA were prepared in single batches. After amplification, cDNA from the entire C1 96-well plate was quantified using picogreen. C1 chips with an average yield of less than 3 nanograms were discarded. The top 15 reactor wells by cDNA concentration were selected as representative 10 pg. samples for sequencing library preparation. Another 50 wells were selected by the same criteria. These were pooled in sets of 10, generating 5 100 pg. samples for each HBR and UHR. All samples for a given source were prepared in a single sequencing library preparation batch using Nextera XT C1 protocol.

#### NuGen

HBR samples were prepared in a single batch using amplification protocol 1, generating 4 10 pg. and 4 100 pg. amplified replicates. UHR samples were prepared in two batches, using either amplification protocol 1 or 2, generating 15 10 pg. and 11 100 pg. samples (see Table S1). A single sequencing library preparation was performed for each batch of samples using either Lucigen NxSeq or NuGen Ovation Rapid protocol (see Table S1).

#### aRNA

Amplification was performed as previously described^26^. HBR samples were prepared in 4 batches from separate dilutions of reference RNA, generating 19 10 pg. and 3 100 pg. amplified replicates. ERCC spike-ins were added to 5 of the 10 pg. replicates before amplification at a dilution of 4,000,000x from stock. UHR samples were diluted and amplified in 2 batches from separate dilutions of reference RNA, generating 12 10 pg. and 7 100 pg. amplified replicates. (Table S1). A single sequencing library preparation was performed using Illumina TruSeq Stranded mRNA protocol modified to begin with amplified aRNA. A small numbers of reads were assigned to ERCC transcripts in replicates from the batch where ERCCs had been added that did not have spike-ins added (average of 0.5% of the number of reads assigned in spiked samples). 18 additional HBR 10 pg. replicates were amplified using aRNA for protocol optimization experiments (see Table S9). These samples were treated separately and were excluded from primary analysis.

#### Bulk UHR and HBR

For each reference RNA, three sequencing libraries were generated from bulk material at the same laboratory as the SmartSeq Plus replicates. Cytoplasmic and mitochondrial ribosomal RNA (rRNA) were depleted using Ribo-Zero Gold as part of Illumina TruSeq Stranded Total RNA protocol. Samples were sequenced on Illumina HiSeq 2000. We also accessed publicly available bulk sequencing of HBR and UHR generated using poly-A selected RNA generated using standard Illumina mRNA-Seq protocol and sequenced on Illumina HiSeq 2000 using 100 bp. paired-end reads. (SEQC/MAQC-III Consortium, 2014,GEO accession numbers: GSM1362002-GSM1362029 (HBR), GSM1361974-GSM1362001 (UHR), downloaded in May 201 5^21^.) These samples were generated as part of a larger experiment to evaluate bulk RNA sequencing where poly-A sequencing was performed at seven sites. For each HBR and UHR, four replicate libraries generated at the NYG site were used. Sequenced read data for each source were pooled. We additionally used publicly available PrimePCR measurements generated by the SEQC/MAQC-III Consortium using UHR and HBR RNA (SEQC/MAQC-III Consortium, 2014, GEO accession number: GPL18522, downloaded in Feb. 2015^21^) to evaluate our reference gene abundance estimates.

Because of differences in experimental design, direct comparison across methods of precision and the effect of input RNA abundance is difficult. For example, input RNA amount as a factor have different meanings for the different amplification methods: for SmartSeq Plus, because 100pg samples were constructed by pooling 10 pg. samples after cDNA amplification, any resulting effects involve library construction, while for aRNA and NuGen resulting effects reflect both cDNA amplification steps and library steps.

### Alignment and quantification

Low confidence nucleotides (with Phred score less than 20) were treated as unknown and replaced with Ns. Unknown nucleotides (Ns) at the ends of reads were trimmed. Poly-A and method-specific adapter sequences were trimmed from the 3’ end of reads using in-house software^27^. Reads were aligned to the human reference genome, build hg19, and to ERCC spike-in transcript sequences using STAR^17^. We retained reads that aligned to at least 40% (paired-end) or 60% (single-end) of trimmed length or 30bp, whichever was greater. In addition, we discarded reads with greater than 30% mismatched positions in trimmed length. Uniquely aligned reads were assigned to GENCODE18 gene annotations and to ERCC transcripts Using HTSeq and htseq-counts. Reads overlapping multiple annotations were assigned to a single gene or discarded using the intersection non-empty method^18^. We normalized raw read counts for differences in sequencing depth using size-factors estimated by the method proposed by Anders and Huber and implemented in DESeq^19^ after filtering genes as described in *Excluded and ambiguous genes,* below. aRNA sequenced data retained RNA strand information, but we did not use this information in quantification so that that all methods were analyzed consistently. For protocol optimization analysis (Table 2), aRNA samples were re-quantified using strand information where applicable. Each method demonstrated different dependence of read counts on gene length (Figure 2H), so no single length normalization procedure was appropriate, hence the analyses were completed without length normalization.

To estimate input RNA abundances, raw sequencing data from all three ribosome-depleted bulk HBR or UHR replicates were pooled resulting in a single sample for each HBR and UHR with sequencing depth of ~400 million reads. Sequencing characteristics of bulk RNA sequencing are relatively well known and we used a model theoretic method to estimate reference gene expression, as implemented in RSEM (RNA-seq by Expectation-Maximization, version 1.2.18, using Bowtie version 1.1.1) strand-specific quantification ^20,28^. Poly-A tails were not added to transcripts. RSEM gene abundances were normalized to transcripts per million (TPM). 50.4% and 51.1% of reads aligned to genes for HBR and UHR, respectively.

We validated the robustness of the RSEM abundance estimates by comparing them to estimates generated using two additional algorithms. First, we used HTSeq and htseq-counts^18^ in the intersection non-empty mode as described above. This method makes few assumptions about the distribution of sequencing reads along transcripts. Second, we used a modified version of Maxcounts^29^, a method designed to be robust to differences in sequencing protocol and each gene was assigned the 95%ile depth of coverage value across covered exons. For both HTSeq and Maxcounts, quantification was strand-specific and estimates were normalized to reads per million (RPM). Counts were also compared to PrimePCR measurements (see *Experimental design).* To compute gene abundance estimates using PrimePCR, we removed undetectable genes (C_T_>35, based on a C_T_ of 35 for one DNA molecule^21^) and then subtracted 35 from each gene's C_T_ value to generate log_2_ number of molecules, which were then converted to log_10_ units. Genes with multiple reported C_T_ measurements were removed, leaving 11,788 (UHR) and 11,572 (HBR) gene measurements for analysis. Pairwise scatter plots and correlations can be found in Figure S1. All quantification algorithms provide similar estimates. We used RSEM quantification throughout because this method provides isoform expression level estimates, which allow more finetuned estimates of gene characteristics (such as GC content and length).

Ribosomal and mitochondrial RNA were depleted from bulk HBR and UHR samples (see *Experimental Design).* We compared estimated RNA abundances based on these samples to abundance based on samples generated using poly-A RNA to determine whether the method of RNA selection substantively affected abundance estimates. Expression estimates were similar across library preparation methods and the library generated with ribosomal and mitochondrial depletion demonstrated the greatest similarity with qPCR measurements (Figure S1B). RSEM expression level estimates based on ribosomal and mitochondrial RNA depleted samples were used as "truth" throughout.

### Excluded and unambiguous genes

We excluded ribosomal genes, genes with short isoforms, and genes on the mitochondrial chromosome, as described in the main text. Inferences made by bioinformatics methods may affect sensitivity, precision, and quantification accuracy for any individual gene. We identified a stringent set of genes to which reads could be uniquely aligned, in order to focus on sensitivity, precision and accuracy of the molecular measurements. Identified genes did not overlap in genomic positions with exons from any other annotated gene on either strand and could be aligned to uniquely across the entire gene. As a measure of mappability we used the GENCODE CRG Alignability track for reference genome hg19, generated by the ENCODE project and downloaded as a bigwig file from the UCSC Genome Browser on Sept. 23 2014^30^. This track contains sliding windows of k-mers and a record of how many locations in the genome each k-mer aligns using the GEM aligner allowing up to two mismatches. We used k equal to 50 nucleotides because the minimum read length in this study was 50 base pairs. Genes where all sliding windows align to only one location were considered uniquely alignable.

### Expected number of molecules in diluted replicate

We estimated the expected number of molecules in a diluted replicate in three steps. First, we estimated the fraction of total input RNA that was targeted for cDNA synthesis and used this to find the mass of targeted RNA. Second, we converted this mass to a total number of input molecules using the average transcript length for each HBR and UHR. Third, we converted gene relative expression levels to expected numbers of molecules in a diluted replicate.

To estimate the mass of RNA targeted for cDNA synthesis, we followed a previously described method^2^. For each SmartSeq Plus dilution replicate, we calculated the percent of reads assigned to ERCC transcripts, with respect to the total number of reads assigned to genes that were retained after filtering. (SmartSeq Plus samples were used because all replicates included ERCC spike-ins.) We divided the known ERCC mass (7.12 or 71.2 femtograms) by the average percentage of reads assigned to ERCC transcripts to get the total mass of targeted transcripts and ERCC molecules and therefore the mass of targeted transcripts. By this method, we estimated the following masses for targeted molecules: 0.24 pg. (HBR 10 pg. replicates), 2.4 pg. (HBR 100 pg. replicates), 0.26 pg. (UHR 10 pg. replicates) and 2.6 pg. (UHR 100 pg. replicates). The average transcript length for HBR, based on RSEM relative gene expression level estimates, was 1,535.56 nucleotides (average transcript mass of 8.175 × 10^-7^ pg. and 288,600 molecules in 10 pg. replicate); for UHR, it was 1,348.39 nucleotides (average mass of 7.179 × 10^-7^ pg. and 364,762 molecules in a 10 pg. replicate). For ERCC molecules, the expected number was calculated directly from the known mass of spiked-in materials and the known molarity of each spike-in transcript. We repeated this analysis using five aRNA HBR 10 pg. samples that contained ERCC spikeins to estimate the mass of targeted mRNA in a diluted HBR replicate. The mass of targeted mRNA was estimated to be 0.15 pg. (HBR 10 pg. replicates). We used RSEM relative gene expression level estimates for poly-A selected bulk HBR samples to estimate the number of targeted mRNA molecules. The average mRNA transcript length in HBR was estimated to be 1,968.73 nucleotides (average mass of 1.048 × 10^-6^ pg. and 143,631 molecules in a 10 pg. replicate).

### Genomic distribution of sequenced reads

Genomic regions were assigned to eight categories hierarchically so that each region was assigned to only one category and so that each read was greedily categorized in the following order: rRNA exon, rRNA repeat, exon (excluding rRNA), intron, flank, intergeneic. Regions were defined based on the following annotations. Exons and introns were assigned based on GENCODE18 annotations. Flanks were assigned to 5 kilobases up‐ and down-stream from gene terminals. rRNA refers to GENCODE18 annotations with "rRNA" as the gene_type, which includes 5S pseudogenes. rRNA repeat refers to RepeatMasker annotations for the rRNA class of repeat. RepeatMasker annotations for reference genome hg19 were downloaded from UCSC table browser as a gtf file from the UCSC genome browser on June 23, 2015. Remaining regions were classified as intergenic. Primary alignments for all reads, including multimapping reads, were assigned to these regions using htseq-counts^31^. The STAR aligner assigns a single primary alignment to each read, with multi-mapping reads assigned the alignment with the best alignment score, if only one such alignment exists, or a randomly selected alignment from the set of best alignments. (Multi-mapping reads were included for this analysis because many rRNA regions demonstrate substantial similarity such that it was difficult to uniquely align reads to these regions.) Haplotype and random chromosomes were excluded.

### Number of detected genes

Genes not observed in the bulk were ignored. The expected number of genes in a diluted replicate was calculated as follows. We assumed that the number of molecules in a tube for a given gene are Poisson distributed with mean equal to the expected number of input molecules and that genes are independent. The presence or absence of a given gene follows a Bernoulli distribution, with the probability of success equal to the probability that at least one molecule for the gene is in the diluted replicate. The number of genes in a diluted replicate is then drawn from a Poisson-Binomial distribution. We used the R package poibin to find a 95% CI for the expected number of genes in a diluted replicate. We performed simulations of the dilution experiment to check robustness of the result to violation of the independence assumption. Simulation results matched theoretical results (data not shown). We performed this analysis both assuming that total RNA was targeted for capture and assuming mRNA was targeted for capture (see *Expected number of molecules in diluted replicates,* above). Because UHR aRNA dilution replicates did not contain ERCC spike-ins, we could only estimate mRNA expectation for HBR.

### Gene traits

We compiled a set of gene characteristics for use in bias exploration. Traits calculated include GC content and length, both known sources of bias for bulk RNA sequencing^32^. Poly-T priming was used by aRNA and SmartSeq Plus and may introduce a bias for genes with internal stretches of adenosines, and so we also computed the presence or absence of an internal A-hexamer (6 or more sequential As). RNA secondary structure may hinder biochemical reactions and we assigned a score for the average strength of local secondary structure. To do this, we calculated the minimum free energy predicted by Vienna RNAFold (version 1.7.2)^33^ for 100 nucleotide-sliding windows along the length of each isoform (step size of 1 nucleotide) and reported the average across all windows. All traits are calculated based on GENCODE18 annotated isoforms. Genes were assigned the average of isoform traits, weighted by the relative expression level of isoforms estimated by RSEM quantification of bulk HBR or UHR. We also calculated two metrics of bioinformatics complexity for each gene. As a measure of alignment complexity, we calculated the fraction of 50 base pair windows that were reported to be uniquely alignable in the GENCODE CRG Alignability track^34^ (see *Excluded and unambiguous genes,* above). As a measure of quantification complexity, we calculated the fraction of the gene body that overlaps with another annotation on either strand. Both of these metrics were calculated over the union of exons for each gene.

### Detection logistic regression

For model fitting, we used computationally unambiguous genes (see *Excluded and unambiguous genes,* above) that were observed in bulk HBR or UHR. Genes within the upper or lower 2.5%ile value for any biophysical trait were excluded so that covariate ranges were well sampled. After filtering, 5,645 genes were included in analysis. The analysis was performed on 10 pg. dilution replicates. 100 pg. dilution replicates were not included because of the small sample size of these groups and because of differences between groups in how these dilution replicates were generated (see Figure 1A and *Experimental design*). A single model was fit containing both HBR and UHR dilution replicates, in order to increase sample size and simplify analysis. A random 90% of the data were used in model development and fitting, with the remaining 10% used to assess model fit. Final sample size for model development was 323,194 observations and for validation it was 45,486 observations.

To determine the best parametric form for each independent variable we followed the multivariate fractional polynomial method. In brief, this method (developed by Royston & Altman, 1994) searches a small range of possible polynomial functions of each independent variable to identify the transform that results in the best model, defined as having the largest log-likelihood. Both one‐ and two-term transforms can be tested. Before selection of a “best” transform, fit models using transformed variables are compared to the linear case (and to each other, if both a one‐ and two-term transformation are considered) using a likelihood ratio test (here the null hypothesis of no difference in fit was rejected at p<0.001). See Hosmer et al.^35^ for more details. In a multivariate case, transformations are tested on individual covariates iteratively in the context of the multivariate model in order of decreasing significance, retaining selected transformations for previously tested covariates. Once all variables have been tested the process repeats, beginning with the previously identified best transforms, until no additional changes are significant. We used a closed test procedure for determining significance (see Hosmer et al.), permitting two-term transformations for the number of molecules and GC content. Single-term transforms were permitted for gene length, strength of local secondary structure and sequencing depth for the sake of model simplicity and interpretability. We used the R mfp package for this analysis^36^. For selecting parametric form, all samples were treated together, ignoring amplification method. By this method, the selected model is:

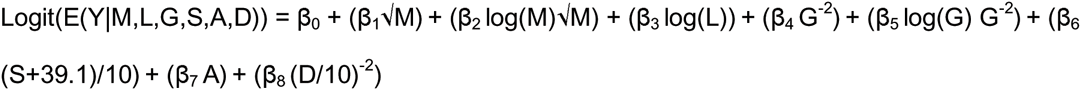

where M represents the expected number of input molecules in a diluted replicate, L represents the gene length (in kilobases), G represents the gene GC content, S represents the gene strength of local secondary structure (kcal/mol, shifted and scaled for stability), A indicates the presence of an A-hexamer within the gene body, and D represents the sequencing depth (per million reads, scaled for stability).

In the final model, amplification method was encoded as dummy variables so that method - specific coefficients were found for all independent covariates, with the exception of sequencing depth. We fit a single coefficient for depth across all methods to increase the covariate range. The final model was fit excluding 17 large influence genes (having Cook’s Distance >0.001 for at least two observations in each of at least two methods) using R built-in glm function with family (error model) set to binomial^37^. The final model can be found in Table S3. Model fit was assessed using normalized Chi-Square (proposed by Osius and Rojek) and normalized Sum-of-Squares goodness-of-fit statistics, evaluated on a random 10% of the data excluded from model development (Table S4, and see Hosmer et al. for details). To assess the benefit of including biophysical and sample covariates, in addition to the expected number of input molecules, we calculated the area under the receiver operating characteristic curve (AUC) for classification using the model, and separately for classification based on the expected number of input molecules alone. AUC provides a measure of the probability that the classifier will assign a higher score to a randomly selected detected gene than a randomly selected undetected gene. AUC average and standard deviations were calculated over 10,000 bootstrap replicates. To determine whether the model was sensitive to read length or paired end status, we calculated fit statistics for data truncated *in silico* to 50 base pair single-end reads (Table S4). We additionally tested extension of model to ERCC spike-in molecules (using SmartSeq Plus and aRNA 10 pg. dilution replicates containing spike-ins) and to dilution samples beginning with 100 pg. input RNA (Table S4). For these additional validations, a random 5,000 observations were used to calculate fit statistics. For tests of extension to 100 pg. data, SmartSeq Plus samples were excluded because these samples were not generated using 100 pg. input RNA for cDNA generation and amplification, but by pooling ten 10 pg. diluted replicates before sequencing library preparation, and so were not appropriate for the modeled process. In all cases, validation statistics were calculated based on predictions for genes within covariate ranges used in model fitting and excluding 17 identified large influence genes. For ERCC samples, this meant that four transcripts shorter than 300 nt. were excluded. Also, because the ERCC molecules span a 10^6^ range while transcriptomes at a single-cell level span ~10^3^ range, 2.5%ile trimming based on input molecules means that only 50 out 92 transcripts were used. While the expected number of input molecules is a very good predictor of gene detection, addition of the remaining independent variables improved prediction (Table S4). All additional independent covariates also contributed significantly to the model. The model was not sensitive to readlength or paired-end status: it fit data truncated *in silico* to 50 base-pair single-end reads well (Table S4). The model did not fit ERCC or 100 pg. dilution replicates well (Normalized Chi-square goodness-of-fit test p < 0.05); however, it still improved prediction accuracy in these cases compared to using the number of input molecules alone for prediction (Table S4).

When examining the effect of the number of input molecules on the probability of gene detection, the remaining covariates were set to median values (gene length of 1.05 kilobases, GC content of 0.49, average strength of local secondary structure of -24.7 kcal/mol, no internal A-hexamer, sequencing depth of 17.1 million reads). To calculate the effect of increasing sequencing depth on percent genes detected, gene detection probabilities were calculated for all genes included in regression analysis (using genespecific covariate values) at each examined depth. The expected number of genes detected is the sum of detection probabilities over all genes. To calculate a molecular detection rate for aRNA with respect to poly-adenylated mRNA molecules, we fit a logistic model with the same functional form using expected number of input molecules calculated from bulk poly-A HBR samples, gene detection data from aRNA HBR 10 pg. dilution replicates, and fixing the depth coefficient to the value estimated in the above analysis.

### Sensitivity outliers

We calculated the squared deviance residual for each observation as a measure of fit, using the logistic model described above. The sum of squared deviance residuals is equivalent to the likelihood ratio test statistic comparing the saturated model with respect to the fitted model, and the sum of squared deviance for a subset of observations can be considered the contribution of this set of observations to overall model fit. To find method-specific problematic genes, we calculated the average squared deviance residual for each gene over all samples for each method separately. For each method, we classified genes with average squared deviance residual larger than 4 as outliers. We repeated outlier identification for computationally ambiguous genes within the range of covariates used in model fitting (n=28,270).

### Coverage

Nucleotide-level coverage was calculated for each gene in the R programming environment^37^ and using Bioconductor libraries GenomicRanges and Rsamtools^38–40^. Coverage was calculated based on uniquely aligned reads only. Only computationally unambiguous genes were used. Additionally, only genes with a single annotated isoform were used in this analysis.

We calculated the observed per nucleotide coverage scaled by the expected coverage as a function of absolute 3' to 5' position within a gene. HBR and UHR dilution replicates were treated together. Replicates were grouped by method and by input amount. Each gene in each sample was considered an independent replicate observation of gene coverage. Genes were filtered to include only those observations with an average of at least 2x coverage per nucleotide. Genes were aligned from the 3’ end, so that the per nucleotide sample size decreased from 3’ to 5’, resulting in increased variance in estimates from 3’ to 5’. Nucleotide positions were filtered to include only those with at least 25 replicate observations, which means that for some genes 5' data was excluded. For each gene, per nucleotide coverage was normalized so that the expected coverage at each position was 1x. For each nucleotide position, the expected value is equal to the number of observations at that position and the observed value is the sum of normalized observed values at that position across observations. Using this this scheme, each gene of at least length *i* contributes equally to the observed coverage at position *i,* regardless of expression level. The result is positional observed / expected coverage values.

To examine patterns of gene coverage as a function of expression level, genes were grouped genes by average per nucleotide coverage. We calculated the average per nucleotide coverage for each of 100 equally sized bins from 5' to 3', rather than coverage as a function of absolute nucleotide position as above, in order to observe qualitative coverage patterns occurring at the same relative position along gene bodies. For each gene, bin values were normalized to sum to one so that within an expression level category all genes contribute equally. For an experimental group, positional bins were assigned the average normalized coverage across all genes, observed in any sample within the experimental group, that fell within a given expression level category.

### Precision

We calculated Pearson pairwise correlation coefficient and Kendall tau pairwise rank correlation coefficient across dilution replicates as a measure of similarity across replicates. The Pearson correlation coefficient is sensitive to large-valued outliers, while the Kendall correlation coefficient is robust. In brief, Kendall correlation is calculated as follows. For each pair of genes the pair is categorized as concordant if the relative ranks of the gene pair are the same for both samples and discordant otherwise. The coefficient reports the fraction of all pairs that are concordant less the fraction that are discordant. For both correlation coefficients, zeros were treated as missing values, such that only genes observed in both members of a pair were included in the calculation.

To characterize measurement precision, we performed least-squares regression of the empirical standard deviation on the empirical mean. We used computationally unambiguous genes to fit these models. Additionally, we included only genes with >95% probability of presence in a diluted replicate, excluded gene detection outliers and trimmed the upper and lower 2.5%ile by mean value for model fitting. Both the average and standard deviation were log-transformed for normality of residuals. After all filtering, at least 1,100 genes were used to fit the model: log_10_(standard deviation) = a + b * log_10_(mean). Sample sizes ranged from 1,149-1,269 genes. A separate model was fit for each experimental group. Because 100 pg. experimental groups have small sample sizes (for most, n<=5) and so provide unstable estimates of variance due to missing values, we performed this analysis on 10 pg. groups only. The NuGen HBR 10 pg. sample size is also quite small (n=4) and was excluded.

To characterize biases in experimental variation we selected a subset of genes where empirical standard deviation was not well predicted by the mean, meaning genes with standardized residuals outside 90% confidence interval of predicted value (assuming a T-distribution with n-3 degrees of freedom for standardized residuals), and a set of "typical" genes, where the gene variance is well predicted by the mean. Typical genes were defined as possessing standardized residuals inside an 80% confidence interval of predicted value. For enrichment tests of GC-content, length, and secondary structure, we calculated the Hodges-Lehmann estimate of difference in location to provide an estimate of the magnitude of in location between test and background gene. This metric estimates the median difference between the two groups.

We identified outliers with unexpectedly high or low experimental variation as genes with 99.3% confidence interval of predicted value. We considered computationally unambiguous genes, and also extended the analysis to computationally ambiguous genes, excluding those with mean expression outside the range used in model fitting.

Principal components analysis was performed on sample covariance matrix calculated using zero-corrected log-transformed read counts for computationally unambiguous genes with non-zero counts in at least on sample and using the R prcomp function. Each PCA included the appropriate bulk HBR or UHR. RSEM-estimated relative frequencies were normalized to the same scale as the diluted replicates using the DESeq method for estimating size factors, as described above. Bivariate normal 95% confidence ellipses were calculated for each experimental group using the R dataEllipse function from the car package^41^.

### Accuracy

Sample sizes (number of genes) for analysis in Figure 4D-N, given filtering described in plot legend, were the following: HBR: n=1,339 (10 pg.) and 2,797 (100 pg.); UHR: n=1,243 (10 pg.) and 2,614 (100 pg.) As stated, in evaluation of gene measurements in individual dilution measurements genes with zero read counts were excluded. For evaluation of average gene measurements, zero values in individual replicates were retained. RSEM-estimated relative frequencies were treated as true relative expression values for each gene. These were normalized to the same scale as the diluted replicates using the DESeq method for estimating size factors, as described above. Wide boxes in boxplots of fold deviation in Figure 4D-F include values for all samples in an experimental group.

To identify method-specific biases in accuracy, we calculated the median fold deviation for each gene across dilution replicates within each experimental group. Genes with fewer than three observations were removed. Of the remaining genes, those with median fold deviation in the upper or lower 5%ile were categorized as overestimated and underestimated, respectively. Remaining genes were used as background for enrichment tests for enrichment. For each method, genes within the upper or lower 1%ile were classified as outlier genes with poor accuracy. Outliers were identified for each experimental group, and then merged across input amounts for each RNA source by taking the union of identified outliers. We repeated outlier identification using computationally ambiguous genes, following the same filtering criteria described above.

To generate density scatter plots of gene read counts in individual dilution replicates, measurements from all 10 pg. dilution replicates for a given method were pooled. The density scatter plots were generated using the R densCols and KernSmooth::bkde2D functions. These functions estimate local density using a binned approximation to a 2 dimensional kernel density with a bivariate Gaussian kernel. log_10_ read counts were used. For density scatter plots of average read counts, averages were taken separately for HBR and UHR 10 pg. dilution replicates. Averages for HBR and UHR were pooled before density calculation.

### Protocol variations

To evaluate the effect of removing purification of initial cDNA, 12 additional HBR 10 pg. dilution replicates were generated. 6 were generated using the same cDNA protocol as the primary aRNA samples, in which initial cDNA is purified using a MinElute column. 6 were generated without this purification step, with adjusted molarity for aRNA amplification to accommodate the change in reaction volume. Each set of 6 included 3 replicates generated using 13 rounds of PCR amplification during sequencing library preparation and 3 using 15. In this analysis, differences in PCR treatment were ignored.

To evaluate the effect of reducing rounds of cDNA amplification, 5 additional HBR 10 pg. dilution replicates were generated using 2 rounds of IVT amplification (rather than 3). All amplified material was used as input for sequencing library preparation. Additionally, these samples were generated without initial cDNA purification and using 15 rounds of PCR during sequencing library preparation (rather than 13). These data were compared to 3 replicates generated using 3 rounds of aRNA amplification, and otherwise following the same protocol. To evaluate an optimized aRNA protocol, excluding initial cDNA purification and reducing rounds of amplification, the same 5 HBR 10 pg. dilution replicates used to examine the effect of reducing rounds of IVT amplification were compared to the primary HBR 10 pg. aRNA data.

To examine the effect of ERCC addition, 10 replicates beginning with 10 pg. HBR total RNA were amplified using aRNA. In 5, ERCC spike-in controls were added with reference RNA at a final dilution of 1:4,000,000. Samples generated in ERCC optimization showed evidence of cross-contamination, with counts assigned to ERCC transcripts (total ERCC counts: 892-1, 457) at appropriate relative abundances for samples generated without addition of ERCC controls.

The effect of strand-specific sequencing was evaluated by re-quantifying aRNA HBR 10 pg. samples using strand information.

## ACKNOWLEDGEMENTS

HD was funded by a National Science Foundation Graduate Research Fellowship; RHC and JAK by the NIH (1U01MH098937); JE and JK by the NIH (5U01MH098953-04); KZ by the NIH (U01MH098977).

## AUTHOR CONTRIBUTIONS

RHC, JE, JAK, KZ and JK designed the study. HD and JK designed and performed computational analysis. RA, AC, BD, RD, OE, JF, JH, TKK, JMK, MYL, RL, WM, JN, JS, NS, TS, AW, CPW, JW, KW, WW and LZ prepared and quality controlled RNA-seq samples, and performed preliminary processing of the sequence data. HD and JK wrote the manuscript.

## MATERIALS & CORRESPONDENCE

Correspondence and material requests should be addressed to Junhyong Kim.

## SUPPLMENETAL FIGURES

**Figure 1:**
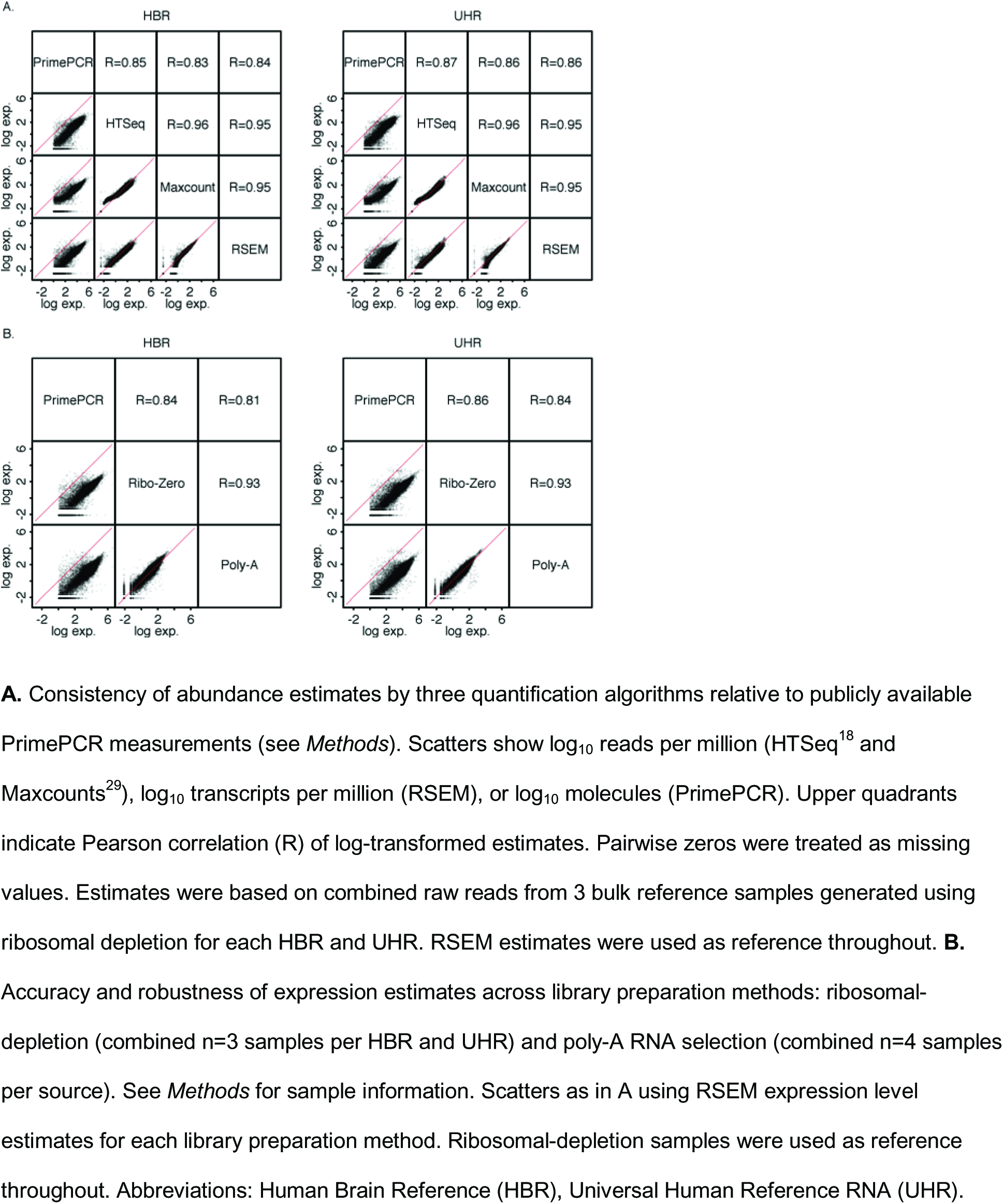
Accuracy and robustness of estimated reference HBR and UHR RNA expression levels

**Figure 2:**
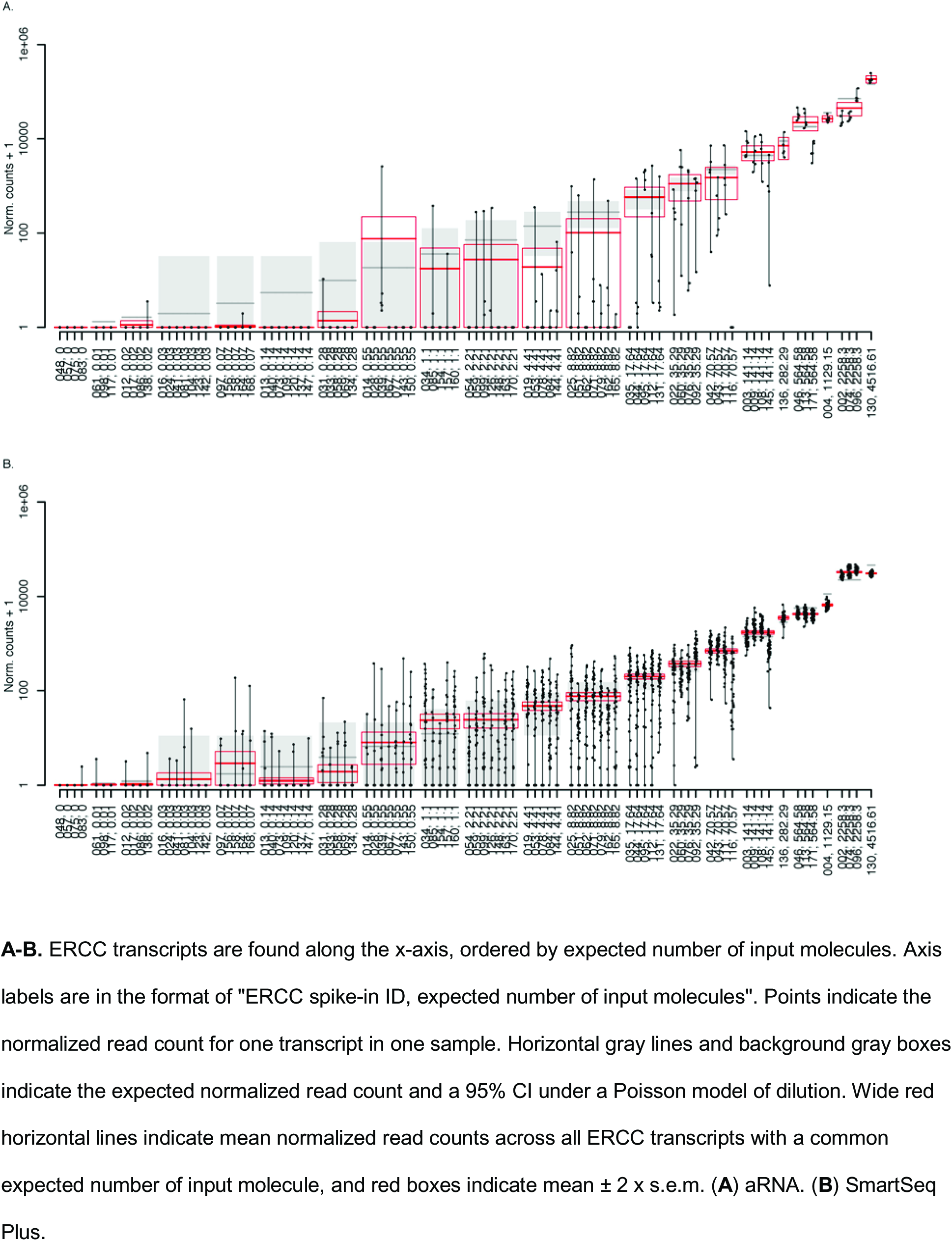
Normalized read counts and expectation for ERCC transcripts

## SUPPLMENETAL TABLES

Supplemental tables can be found in attached “supplementalTables.xlsx” file.

**Table S1.** Control dataset sample identification, protocol information, and RNA sequencing stats Experimental group, protocol information and RNA sequencing statistics for each sample used in primary analyses. Alignment statistics were based on STAR alignment to hg19 and were with respect to reads retained after trimming for primer or poly-A sequences^17^.

**Table S2.** Computationally unambiguous genes

Genes to which reads can be uniquely assigned. See the <u>*Excluded and unambiguous genes* section in</u> <u>*Methods* for details on classification.</u>

**Table S3.** Gene detection logistic regression model

See model details in *Methods. Abbreviations:* M expected number of input molecules; L gene length (kilobases); G gene GC content; S strength of gene local secondary structure (kilocalories per mole); hasA presence of A-hexamer internal to gene body; D Depth (per 10,000,000 reads); *S.E.* standard error; *Wald Z* Wald test statistic; *Pr*(>*Z*) Wald test p-value.

**Table S4.** Gene detection logistic regression fit and validation

Model was fit using randomly selected 90% of 10 pg. data, excluding 17 large influence genes. Fit was evaluated on the remaining 10% of the data. Fit was also evaluated on sequence data that was *in silico* truncated to 50 base pair single end (“Truncated”), ERCC read counts (“ERCC”), and 100 pg. dilution replicates (“100 pg.”). AUC (area under receiver operating characteristic curve) reported as mean values ± 2 Sd. calculated over 10,000 bootstrap samples. AUC (molecules) predicts detection based on number of input molecules alone. See *Methods* for further details.

**Table S5.** Probability of gene detection

Based on model described in *Methods.* Remaining covariates set to median value.

**Table S6.** Gene detection outliers

Genes that are problematic for detection. See *Methods* for classification of outliers. "Gene set" indicates whether gene is classified as computationally unambiguous (1) or not (2). "Detected / undetected" indicates whether the gene is unexpectedly observed (D) or unexpectedly unobserved (U).

**Table S7.** Precision outliers

Genes with standardized residual outside a 99.3% confidence interval, with respect to regression of standard deviation on the mean (see *Methods).* "Gene set" indicates whether gene is classified as computationally unambiguous (1) or not (2). Only genes whose mean is within the range of fitted model were included. Column values indicate whether indicate whether the gene standard deviation is unexpectedly low (L) or high (H), given mean.

**Table S8.** Accuracy outliers

Genes were identified as accuracy outliers if its median fold deviation, taken across dilution replicates, was contained in the upper or lower 1%ile of all considered genes (see *Methods).* Columns labeled by single-cell protocol contain an "H" if a gene was identified as an overestimated outlier, and an “L” if a gene was identified as an underestimated outlier. "Gene set" indicates whether gene is classified as computationally unambiguous (1) or not (2).

**Table S9.** Optimization dataset sample identification, protocol information, and RNA sequencing stats As Table S1 for samples used in protocol optimization analyses.

